# Functional interpretation of *ATAD3A* variants in neuro-mitochondrial phenotypes

**DOI:** 10.1101/2020.10.05.318519

**Authors:** Zheng Yie Yap, YoHan Park, Saskia B. Wortmann, Adam C. Gunning, Sukyoung Lee, Lita Duraine, Ekkehard Wilichowski, Kate Wilson, Johannes A. Mayr, Matias Wagner, Hong Li, Usha Kini, Emily Davis Black, James R. Lupski, Sian Ellard, Dominik S. Westphal, Tamar Harel, Wan Hee Yoon

**Affiliations:** Aging and Metabolism Research Program, Oklahoma Medical Research Foundation, Oklahoma City, OK, USA; Institute of Human Genetics, Technical University Munich, Munich, Germany; University Children’s Hospital, Paracelsus Medical University (PMU), Salzburg, Austria; Radboud Centre for Mitochondrial Medicine (RCMM), Amalia Children’s Hospital, Nijmegen, The Netherlands; Exeter Genomics Laboratory, Royal Devon and Exeter NHS Foundation Trust, Exeter EX2 5DW, UK; Institute of Biomedical and Clinical Science, College of Medicine and Health, University of Exeter, Exeter EX2 5DW, UK; Verna and Marrs McLean Department of Biochemistry and Molecular Biology, Baylor College of Medicine, Houston, TX, USA; Jan and Dan Duncan Neurological Research Institute, Baylor College of Medicine, Houston, TX, USA; Department of Pediatrics and Pediatric Neurology, University Medical Center Göttingen, Georg-August-Universität Göttingen, Göttingen, Germany; Oxford Centre for Genomic Medicine, Oxford University Hospitals NHS Foundation Trust, Oxford, UK; Institute of Neurogenomics, Helmholtz Zentrum München, Neuherberg, Germany; Department of Human Genetics, School of Medicine, Emory University, Atlanta, GA, United States; Department of Pediatrics, School of Medicine, Emory University, and Children’s Healthcare of Atlanta, Atlanta, GA, United States; Department of Molecular and Human Genetics, Baylor College of Medicine, Houston, TX 77030, USA; Department of Pediatrics, Baylor College of Medicine, Houston, TX 77030, USA; Human Genome Sequencing Center, Baylor College of Medicine, Houston, TX 77030, USA; Texas Children’s Hospital, Houston, TX 77030, USA; Department of Genetics, Hadassah-Hebrew University Medical Center, Jerusalem, Israel 9112001

**Keywords:** ATAD3A, mitochondria, disease, autosomal recessive, autophagy, neurogenesis, Drosophila, AAA+ protein

## Abstract

**Background:** The ATPase family AAA-domain containing protein 3A (ATAD3A) is a nuclear-encoded mitochondrial membrane anchored protein involved in diverse processes including mitochondrial dynamics, mitochondrial DNA organization, and cholesterol metabolism. Biallelic deletions (null), recessive missense variants (hypomorph), and heterozygous missense variants or duplications (antimorph) in *ATAD3A* lead to neurological syndromes in humans.

**Objective:** To expand the mutational spectrum of *ATAD3A* variants and to provide functional interpretation of missense alleles in trans to deletion alleles.

**Methods:** Exome sequencing was used to identify single nucleotide variants (SNVs) and copy number variants (CNVs) in *ATAD3A* in individuals with neurological and mitochondrial phenotypes. A Drosophila Atad3A Gal4 trap null allele was generated using CRISPR-Cas9 genome editing technology to aid interpretation of variants.

**Results:** We report 13 individuals from 8 unrelated families with biallelic *ATAD3A* variants. Four of the identified missense variants, p.(Leu77Val), p.(Phe50Leu), p.(Arg170Trp), p.(Gly236Val), were inherited in trans to loss-of-function alleles. A fifth missense variant, p.(Arg327Pro), was homozygous. Affected individuals exhibited findings previously associated with *ATAD3A* pathogenic variation, including developmental delay, hypotonia, congenital cataracts, hypertrophic cardiomyopathy, and cerebellar atrophy. Drosophila studies indicated that Phe50Leu, Gly236Val, and Arg327Pro are severe loss-of-function alleles leading to early developmental lethality and neurogenesis defects, whereas Leu77Val and Arg170Trp are partial loss of function alleles that cause progressive locomotion defects. Moreover, Leu77Val and Arg170Trp expression leads to an increase in autophagy and mitophagy in adult muscles.

**Conclusion:** Our findings expand the allelic spectrum of ATAD3A variants, and exemplify the use of a functional assay in Drosophila to aid variant interpretation.

## BACKGROUND

The ATPase family AAA-domain containing protein 3A (ATAD3A) belongs to a family of hexameric ATPases associated with diverse cellular activities (AAA+ ATPase proteins). It was initially identified as a mitochondrial protein enriched at contact sites between the mitochondria and the endoplasmic reticulum (ER) membrane.^1^ The protein is presumed to tether the inner mitochondrial membrane to the outer mitochondrial membrane and has the capacity to interact with the ER, thus potentially regulating mitochondria-ER interorganellar interactions and exchanges.^1; 2^ Several studies have alluded to the importance of ATAD3A in embryonic development. Deletion of *Atad3a* in mice causes embryonic lethality at day E7.5, with growth retardation and defective development of the trophoblast lineage.^3^ Knockdown of the *Drosophila* ortholog, *belphegor*, (*bor, dAtad3a*) results in growth arrest during larval development,^1^ and the *C. elegans* ortholog is essential for mitochondrial activity and development.^4^ RNAi studies of human *ATAD3A* in lung cancer cells have documented increased mitochondrial fragmentation and a decreased co-localization of mitochondria and endoplasmic reticulum (ER).^5^

In humans, the ATAD3 gene family contains three paralogs that appear to have recently evolved by duplication of a single ancestral gene: *ATAD3A, ATAD3B*, and *ATAD3C*. These are located in tandem and map to chromosome 1p36.33.^3; 6^ The major *ATAD3A* isoform, p66, is ubiquitously expressed, whereas the major *ATAD3B* isoform, p67, is specifically expressed during development^6^ and reactivated in cancer.^7; 8^ *ATAD3C* lacks four exons, suggesting that it may be a pseudogene. The genetic architecture dictated by three highly homologous paralogs predisposes the region to genomic instability and rearrangements generated by nonallelic homologous recombination (NAHR).^9; 10^

To date, the allelic spectrum of *ATAD3A-*associated disease [MIM: 617183] includes null, hypomorph, and antimorph alleles.^10-14^ Biallelic deletions mediated by NAHR, most often spanning ∼38kb between *ATAD3B* and *ATAD3A* and less frequently ∼67kb between *ATAD3C* and *ATAD3A*, lead to an infantile-lethal presentation including respiratory insufficiency, neonatal seizures, congenital contractures, corneal clouding and/or edema, pontocerebellar hypoplasia and simplified sulcation and gyration.^12^ Deletions between *ATAD3B* and *ATAD3A* lead to a fusion transcript under regulation of the weaker *ATAD3B* promoter, and thus show decreased expression of an ATAD3B/ATAD3A fusion protein that presumably is sufficient for fetal development but apparently cannot sustain life beyond the neonatal period.^12^ The reciprocal, NAHR-mediated duplication at this locus, between *ATAD3C* and *ATAD3A*, results in a fusion gene encoding a dysfunctional protein.^15^ Homozygosity for presumed hypomorphic missense alleles (p.Thr53Ile, p.Thr84Met) leads to bilateral cataracts, hypotonia, ataxia and cerebellar atrophy.^10; 16^ Finally, a recurrent *de novo* heterozygous missense variant (p.Arg528Trp) acts as an antimorph or dominant-negative allele and gives rise to a phenotypic spectrum including developmental delay, hypotonia, optic atrophy, axonal neuropathy, and hypertrophic cardiomyopathy.^10^

We report on the clinical and molecular findings of 13 individuals from 8 families with biallelic variants at the *ATAD3A* locus, and expand the allelic spectrum to include those with a missense variant inherited in *trans* to an expected loss-of-function (deletion or frameshift) allele. *In vivo* functional studies for the missense variants in *ATAD3A* using a *Drosophila* model revealed that these were hypomorphic alleles that exhibited diverse allelic strength, and shed light on genotype-phenotype correlations.

## METHODS

### Exome analysis

Following informed consent, exome sequencing was pursued on DNA extracted from whole blood of affected individuals from each of 8 unrelated families. Study design was adapted to each family, and was either proband-only, trio (parents and affected child), or sibship analysis. For Families 2, 4, 5, and 6, sample collection, DNA extraction, exome library preparation, sequencing, and variant calling and annotation were performed as previously described.^17^ For Family 6, trio whole exome sequencing was performed using the proband’s and unaffected parents’ samples. Exome read-depth was assessed manually. For Families 1, 7, and 8, exome sequencing and data analysis were as previously described in Wagner et al. (2019).^18^

### Sanger validation and segregation of the variants

Single nucleotide variants (SNV) of interest were confirmed by Sanger sequencing, and segregation of variants was carried out in available family members. Intergenic *ATAD3B-ATAD3A* copy number variants were not confirmed, as these have been well established in recent literature and exome read-depth data was compelling.

### 3D Modeling of Protein Structure

The 3D model was predicted with I-TASSER.^19^ After inspection of the predicted model, two helices (residues 225-242 and 247-264) were inserted manually. Spatial arrangement of secondary structures was adjusted manually to ensure proper domain separation by the mitochondria inner membrane.

### Cloning and Transgenesis

*dAtad3a-T2A-Gal4* allele was generated using modified methods of CRISPR/Cas-9-mediated genome editing^20^ and homology-dependent repair^21^ by WellGenetics Inc. We targeted the first coding intron of *dAtad3a* using gRNAs (TGTGATAGCGTGGCGCATGC[CGG]). The gRNA was cloned into an U6 promoter plasmid. Cassette T-GEM(1) is composed of an attP site, a linker for phase 1 in-frame expression, T2A, Gal4, Hsp70 transcription terminator, a floxed 3xP3-RFP, and an inverted attP (*attB-splicing acceptor-T2A-Gal4-polyA-loxP-3xP3-RFP-loxP-attB*).^22^ The cassette and two homology arms were cloned into pUC57-Kan as a donor template for repair. The cassette contains an upstream homology arm (HA_L_ - 1016bp) and a downstream homology arm (HA_L_ - 1039bp). The homology arms were amplified using primers: HA_L_ -F: 5’-GCACGCCCACAATTAGCATT-3’, HA_L_-R 5’-GGTTATGCAATTGGCTGATGAAA-3’, HA_R_-F: 5’-GGAGGCCCTCGAGCTGTC -3’, HA_R-_R: 5’-CCAGTCGAACACCTTGTGGA - 3’. The gRNA and hs-Cas9 were supplied in DNA plasmids, together with donor plasmid for microinjection into embryos of control strain *w*^*1118*^. F1 progenies carrying selection marker of 3xP3-RFP were further validated by genomic PCR and Sanger sequencing. 3xP3-RFP was removed by Cre recombinase.

For construction of pUASTattB-dAtad3a^WT^-V5, a full-length dAtad3a cDNA was amplified by PCR from a pUAST-dAtad3a clone^10^, and then subcloned into a pUASTattB vector^23^ using primers: 5’-GGATCCaaa**ATG**TCGTGGCTTTTGGGCAGG -3’, 5’-GCGGCCGC**TTA**GGTGCTATCCAGTCCGAGCAGTGGATTCGGGATCGGCTTGCCGCC GCTTCC CAGTTTCTTTGCAGTTAGGGTG-3’.

pUASTattB-dAtad3a^L83V^-V5, pUASTattB-dAtad3a^F56L^-V5, pUASTattB-dAtad3a^R176W^-V5, pUASTattB-dAtad3a^G242V^-V5, and pUASTattB-dAtad3a^R333P^-V5 were generated by site-directed mutagenesis PCR using primers: (L83V)-F: 5’-cacgcccgggaggccctcgagGTGtccaagatgcaggaggccacc -3’, (L83V)-R: 5’-ggtggcctcctgcatcttggaCACctcgagggcctcccgggcgtg-3’, (F56L)-F: 5’-aaggccatggaagcgtaccgcTTAgatTCGTCGGCGCTGGAACGT-3’, (F56L)-R: 5’-ACGTTCCAGCGCCGACGAatcTAAgcggtacgcttccatggcctt-3’, (R176W)-F: 5’-gtccagcgtcaagaggccatgTGGcgccagaccatcgagcacgag -3’, (R176W)-R: 5’-ctcgtgctcgatggtctggcgCCAcatggcctcttgacgctggac -3’, (G242V)-F: 5’-gctggtactgttatcggtgccGtTgctgaggctatgcttaccgac-3’, (G242V)-R: 5’-gtcggtaagcatagcctcagcAaCggcaccgataacagtaccagc-3’, (R333P)-F: 5’-ctaaatccgaagctggaggaaCcGcttcgtgacattgccatcgcc-3’, (R333P)-R: 5’-ggcgatggcaatgtcacgaagCgGttcctccagcttcggatttag-3’. A series of pUASTattB-dAtad3a constructs were injected into *y,w,ΦC31; VK37* embryos, and transgenic flies were selected.

### Fly Strains and Maintenance

The following stocks were obtained from the Bloomington Stock Center at Indiana University (BDSC): *w*^*1118*^; *PBac{PB}bor*^*c05496*^*/TM6B, Tb*^*1*^, *w*^*1118*^; *Df(3R)Excel7329/TM6B, Tb*^*1*^, and *w*; 20xUAS-IVS-mCD8::GFP* (on III). All flies were maintained at room temperature (21°C). All crosses were kept at 25°C.

### Western Blotting

Fly heads were homogenized in 1x Laemmli sample buffer containing 2.5% β-mercaptoethanol (Sigma-Aldrich). After boiling for 10 min, samples were loaded into 4–20% Mini-PROTEAN® TGX Stain-Free(tm) Protein Gels (Bio-Rad), separated by SDS-PAGE, and transferred to nitrocellulose membranes (Bio-Rad). The primary antibodies were used for overnight shaking at 4°C by the following dilution: mouse anti-V5 (Invitrogen Cat# R960-25 RRID: AB_2556564), 1:2,000; mouse anti-ATP5A (Abcam Cat# ab14748 RRID: AB_301447), 1:2000; mouse anti-Actin (MP Biomedicals Cat# 8691002), 1:20,000; rabbit anti-Ref2(P) (kindly provided by Sheng Zhang). HRP conjugated goat anti-rabbit (Invitrogen Cat# G-21234 RRID: AB_2536530), anti-mouse (Invitrogen Cat# A-28177 RRID: AB_2536163) were used at 1:7,000, and visualized with ECL(Bio-Rad).

### Embryo Collection and Immunostaining

Embryos were collected on grape juice plates for 24 hours at 37°C. Collected embryos were washed twice with deionized water, and dechorionated with 50 % bleach for 3 minutes. After rinsed thoroughly, embryos were fixed for 30 minutes by 1:1 ratio of Heptane (Sigma Aldrich Cat #246654-1L) and 4 % formaldehyde (Thermo Fisher Cat#F79500) in 1 x Phosphate Buffered Saline (PBS), pH 7.4. To remove vitelline membranes, embryos were shaken vigorously in methanol for 5 times. 1 x PBS, pH 7.4 containing 0.2% BSA and 0.3 % Triton-X100 were used for rehydration and washing. The primary antibodies were used for overnight at the following dilutions: rat anti-Elav 1:500 (DSHB Cat# 7E8A10 RRID:AB_528218), rabbit anti-β-galactosidase 1:250 (Invitrogen Cat# A-11132 RRID: AB_221539), rabbit anti-GFP 1:1000 (Invitrogen Cat# A-11122 RRID:AB_221569), Alexa 647 conjugated goat anti-Horseradish Peroxidase 1:500 (Jackson ImmunoResearch Labs Cat# 123-605-021 RRID: AB_2338967). Alexa 488 conjugated anti-rat (Invitrogen Cat# A-21208 RRID: AB_2535794), and Alexa 568 conjugated anti-rabbit (Invitrogen Cat# A-11011 RRID: AB_143157) secondary antibodies were used at 1:500. Samples were mounted in Vectashield (Vector Labs Cat# 10198-042, Burlingame, CA). Imaging was performed using LSM710 confocal microscope (Zeiss). Images were processed with Zeiss LSM Image Browser and Adobe Photoshop.

### Adult Drosophila Thorax Sectioning

Flies were fixed in 4% formaldehyde with PBS containing 0.3% TritonX-100 for 4 hours at 4’C on rotator and rinsed with PBS to remove any residual formaldehyde. Fixed flies were then dissected in PBS. Firstly, fly wings were remove carefully without tearing the tissue of the thorax. Then the head and abdomen together with the intestines are removed so only the thorax would remain. Holding onto the legs as support to stabilize the thorax, a sharp blade was used make a slice down the middle of the thorax (dorsal side). The thorax was transferred into an Eppendorf tube and washed with PBS to remove any debris. The primary antibodies were used at the following dilutions: mouse anti-ATP5A 1:500, rabbit anti-Ref2p 1:1000. Alexa 488 conjugated and Alexa 568 conjugated secondary antibodies were used at 1:500. Samples were mounted in Vectashield. Imaging was performed using LSM710 confocal microscope (Zeiss). Images were processed with Zeiss LSM Image Browser and Adobe Photoshop.

### Drosophila Flight Assay

The method was adapted from Pesah et al. (2004).^24^ Flies were anesthetized and allocated into individual food vials for 24 hours at 25°C before the assays were performed to allow full recovery from the effects of CO_2_. Each individual vial was inverted into a 500mL measuring cylinder and gently taped to dislodge the fly into the cylinder. Flies either fell to the bottom in a straight line or flew to the side of the cylinder. Recording of the whole process was taken and analysis is done based on each behavior the flies exhibited. 25 flies of each genotype were assayed.

### Drosophila Climbing Assay

Method was adapted from Madabattula et al. (2015).^25^ 25 flies were anesthetized using CO_2_ and allowed to rest in fresh food vials 24 hours at 25°C prior to the assay. Male and female flies were kept separately as gender difference on behavior might be significant. To prepare the climbing apparatus, measure a distance of 8 cm from the bottom surface of an empty polystyrene vial and mark the distance by drawing a line around the entire circumference of the vial. Flies were transferred without using CO_2_ into different climbing apparatus for each genotype to prevent cross contamination. The apparatus was closed off by vertically joining to another empty polystyrene vial using tape and the flies were left to acclimatize to the surrounding for at least 10 min. Then, the apparatus was gently tapped five times to displace the flies to the bottom of the apparatus and a video was recorded for 20 s to measure the number of flies able to cross the height of 8 cm at each time point. After a 10 min rest, the assay was repeated. Three trials were conducted.

### Transmission Electron Microscopy

Drosophila thoraxes were imaged following standard Electron Microscopy procedures using a Ted Pella Bio Wave processing microwave with vacuum attachments. Briefly, whole thorax with wings were dissected at room temperature in modified Karnovski’s fixative in 0.1 M Sodium Cacodylate buffer at pH 7.2 and subsequently fixed overnight to three days in the same fixative. The pre-fixed thoraxes were then irradiated and fixed again, followed by 3x millipore water rinses, post-fixed with 1% aqueous Osmium Tetroxide, and 1% Potassium Ferrocyanide mixture in Millipore water. This was followed by 3X Millipore water rinses. Ethanol concentrations from 30-100% were used for the initial dehydration series, followed with 100% Propylene Oxide as the final dehydrant. Samples were gradually infiltrated with 3 ratios of propylene oxide and Embed 812, finally going into 3 changes of pure resin under vacuum. Samples were allowed to infiltrate in pure resin overnight on a rotator. The samples were embedded into flat silicone molds and cured in the oven at 62°C for at least three days. The polymerized samples were thin-sectioned at 48-50 nm and stained with 1% uranyl acetate for thirteen minutes followed by 2.5% lead citrate for two and a half minutes the day before TEM examination. Grids were viewed in a JEOL 1400 Plus transmission electron microscope at 80kV. Images were captured using an AMT XR-16 mid-mount 16 mega-pixel digital camera in Sigma mode. Images were contrast adjusted in Image J.

## RESULTS

### Clinical Reports

Biallelic variants in *ATAD3A* were identified in 13 individuals from 8 previously unreported families (**Figure 1A**). These included different combinations of copy number variants (CNV) and/or single nucleotide variants (SNV). Detailed clinical presentations are supplied in **Table 1 and Table S1 (Clinical information -Additional file 1)**. Briefly, individuals with biallelic deletion CNVs showed a neonatal-lethal presentation with respiratory failure, generalized hypotonia, seizures, congenital contractures, bilateral ophthalmologic findings, and brain anomalies, consistent with the reported phenotype.^12^ Individuals with a loss-of-function allele (intergenic CNV, intragenic CNV, or frameshift SNV) inherited in *trans* to a missense variant (Families 3-6) presented with varied severity of the phenotype (**Table 1, and Table S1 - Clinical information -Additional file 1**), which we hypothesized to correlate with the degree of pathogenicity of the missense alleles. Finally, two families with other combinations of biallelic variants (nonframeshift indels or missense variants, Families 7-8) presented with *ATAD3A-* associated features such as cataracts and hypertrophic cardiomyopathy. Affected individuals in Family 8 exhibited severe phenotypes and lethality at 6-7 months after birth. Variants identified by exome data analysis were tested for segregation with the disease by Sanger sequencing. Of note, in Family 7, the c.150C>G; p.(Phe50Leu) was *de novo* in the proband, and presumably arose on the paternal allele, since the c.1703_1705delAGA variant was inherited from the mother.

**Table 1.** Summary of significant clinical findings

|  | Family 1 | Family 2 | Family 3 | Family 4 (2 siblings) | Family 5 (3 siblings) | Family 6 | Family 7 | Family 8 (3 siblings) |
| --- | --- | --- | --- | --- | --- | --- | --- | --- |
| **ATAD3A variants (NM_001170535.1)** | *ATAD3B-ATAD3A* ~38KB deletion (hom) | *ATAD3B-ATAD3A* comp het deletion | *ATAD3B-ATAD3A* del;<br>c.229C>G;<br>p.(Leu77Val) | Exon 3-4 del (c.(282+1_283-1)_(444+1_445-1)del<br>c.150C>G;<br>p.(Phe50Leu) | c.1141dup;<br>p.(Val381Glyfs*17)<br>c.508C>T,<br>p.(Arg170Trp) | c.1414del<br>p.(His472fs)<br>c.707G>T,<br>p.(Gly236Val) | c.150C>G,<br>p.(Phe50Leu);<br>c.1703_1705delAG<br>A, p.(Lys568del) | c.980G>C,<br>p.(Arg327Pro)<br>(hom) |
| **Age at last exam** | 13 d (deceased) | 30 hours (deceased) | 19 mo (deceased at 2 y) | Died shortly after birth | 17-19 years | 3 mo | 15 y | 6-7 mo (all deceased) |
| **Developmental delay** | NR | NR | global DD | NR | moderate-severe learning difficulties | NA | mild DD | NR |
| **Neurological exam** | hypotonia, no respiratory effort | hypotonia, no respiratory effort | NA | hypotonia, no respiratory effort | ataxia, muscle wasting | central hypotonia, increased peripheral tone | mild hypotonia, | NA |
| **Congenital cataract** | + | cloudy corneas | + | NA | one of 3 siblings | - | + | + |
| **Hypertrophic cardiomyopathy** | + | + | + | + | - | + | + | + |
| **Cerebellar atrophy/hypoplasia** | + | + | NA | + | + | + | + | NA |
| **Elevated 3-methylglutaconate in urine** | NA | NA | NA | + | NA | NA | NA | + |
| **Other** | - | undescended testes | - | - | hearing loss | - | GH deficiency | - |

Abbreviations: DD – developmental delay; dup – duplication; GH – growth hormone; hom – homozygous; NA – not available

**Figure 1.**
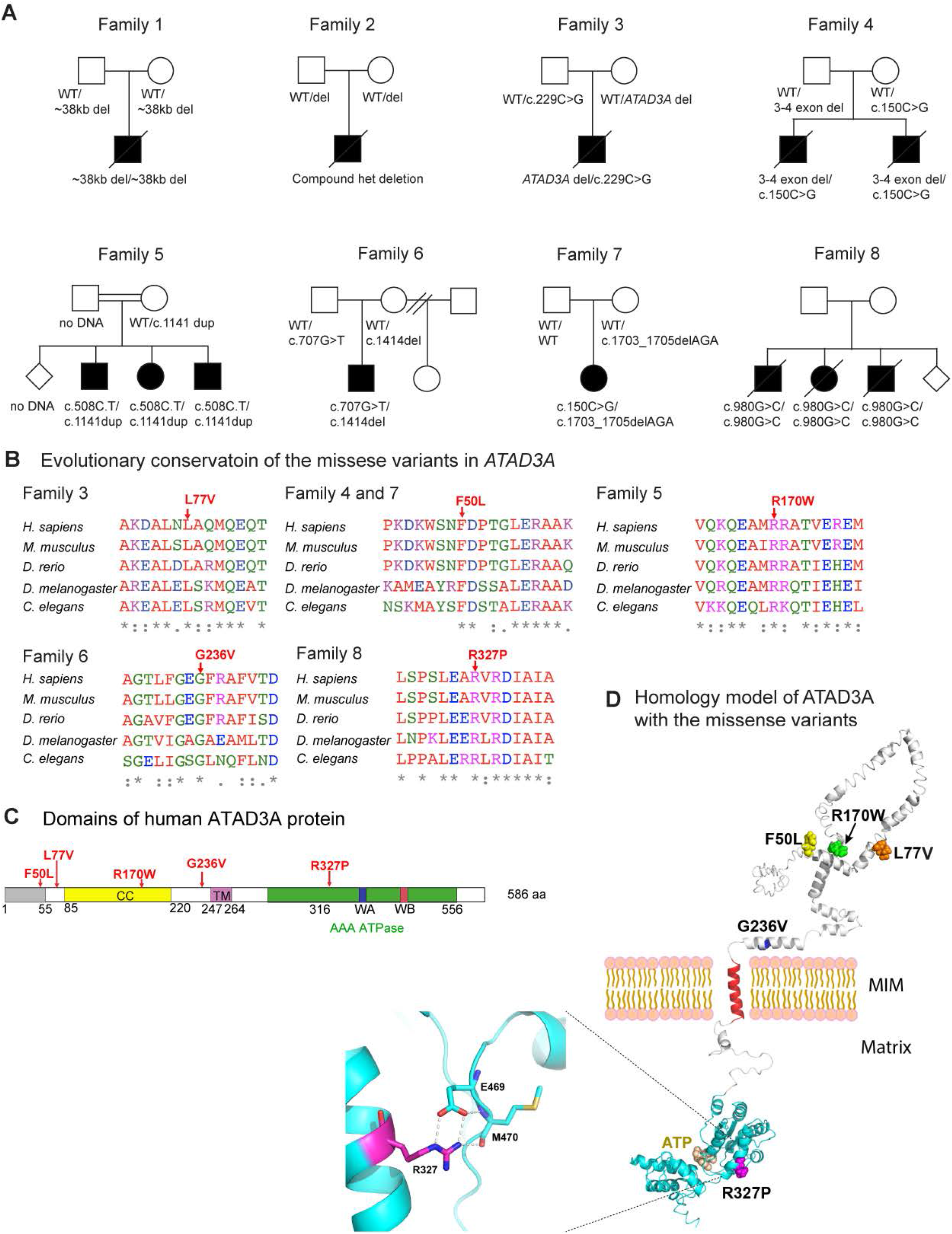
Identification of Patients with Neurological Phenotypes with Variants in *ATAD3A*. (A) Pedigrees of studied families, indicating biallelic variants in *ATAD3A* identified in 13 individuals from 8 families, indicating biallelic deletion in family 1 and 2, loss-of-function alleles (intergenic CNV, intragenic CNV, or frameshift SNV) inherited in *trans* to a missense variant in Families 3-7, homozygous missense variant in Family 8. (B) Protein sequence alignment in multiple species confirms evolutionary conservation of p.L77V, p.F50L, pR170W, p.G236V, p.R327P, in both humans and Drosophila. (C) Schematic representation of protein domains of human ATAD3A. CC indicates coiled-coil domain. TM indicates putative transmembrane domain. Green indicate AAA+ domain containing Walker A motif (WA) and Walker B motif (WB). (D) In silico protein structure prediction of ATAD3A shows position of mutated residues.

### *In silico* analysis of missense variants identified in affected individuals

The five *ATAD3A* missense variants identified in affected individuals include (provided according to NM_001170535.3, see **Table 2**): c.150C>G, p.(Phe50Leu), c.229C>G, p.(Leu77Val), c.508C>T, p.(Arg170Trp), c.707G>T, p.(Gly236Val), and c.980G>C, p.(Arg327Pro). These variants will be referred to as F50L, L77V, R170W, G236V, and R327P, respectively. The variants have not been reported previously as pathogenic variants and were not seen in the homozygous state in gnomAD, the largest available population database (https://gnomad.broadinstitute.org/) (https://www.biorxiv.org/content/10.1101/531210v3). All of the altered residues are evolutionarily conserved in species of the animal kingdom (Figure 1B).

**Table 2.**
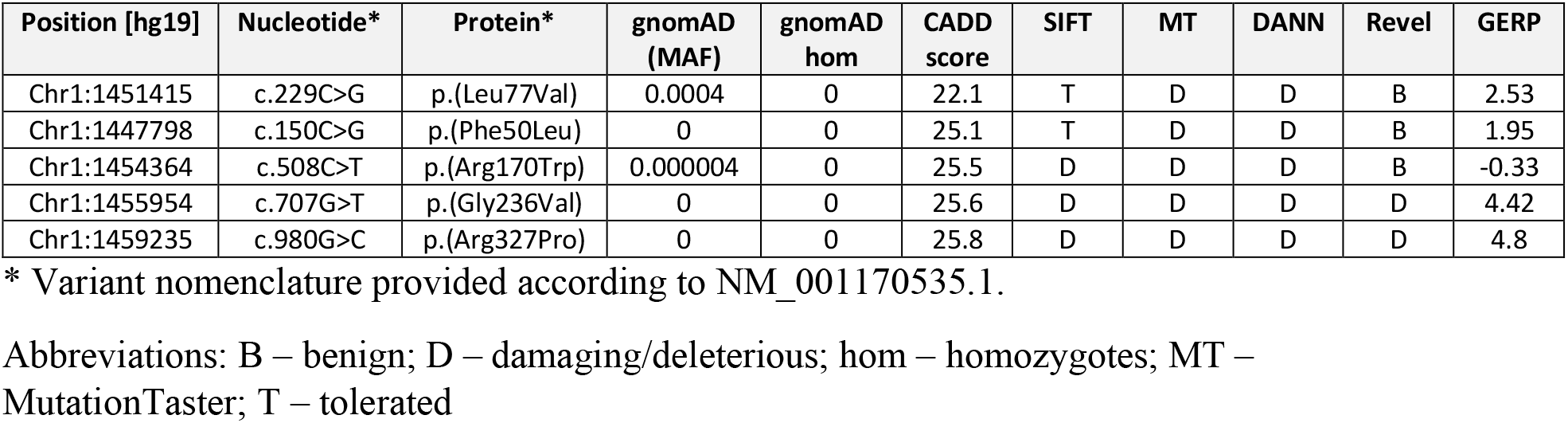
Missense variants identified in *ATAD3A*

| Position [hg19] | Nucleotide* | Protein* | gnomAD (MAF) | gnomAD hom | CADD score | SIFT | MT | DANN | Revel | GERP |
| --- | --- | --- | --- | --- | --- | --- | --- | --- | --- | --- |
| Chr1:1451415 | c.229C>G | p.(Leu77Val) | 0.0004 | 0 | 22.1 | T | D | D | B | 2.53 |
| Chr1:1447798 | c.150C>G | p.(Phe50Leu) | 0 | 0 | 25.1 | T | D | D | B | 1.95 |
| Chr1:1454364 | c.508C>T | p.(Arg170Trp) | 0.000004 | 0 | 25.5 | D | D | D | B | -0.33 |
| Chr1:1455954 | c.707G>T | p.(Gly236Val) | 0 | 0 | 25.6 | D | D | D | D | 4.42 |
| Chr1:1459235 | c.980G>C | p.(Arg327Pro) | 0 | 0 | 25.8 | D | D | D | D | 4.8 |

\* Variant nomenclature provided according to NM\_001170535.1.
Abbreviations: B – benign; D – damaging/deleterious; hom – homozygotes; MT – MutationTaster; T – tolerated

In silico structural modeling of ATAD3A by PredictProtein^26^ suggested that each amino acid variant would alter the predicted protein structure (Figure 1D). Phe50 is located next to the first α-helix following the disordered region. Leu77, Arg170, Gly236 and Arg327 are strictly conserved and found in α-helices. Leu77 is predicted to be buried inside the ATAD3A structure and is not exposed at the surface. The p.(Leu77Val) variant could affect a hydrophobic interaction between the α-helix and other part of the structure due to shortening of the side chain by one carbon. On the contrary, Arg170 is predicted to be surface exposed. Hence, the p.(Arg170Trp) variant may increase surface hydrophobicity, reducing solubility and resulting in a less stable protein. Gly236 is located next to the GxxFG motif that guides the folding and assembly of the membrane-spanning amphipathic α-helix. The p.(Gly236Val) variant may affect the interaction between this potential transmembrane helix and the mitochondrial inner or outer membrane or between neighboring transmembrane helices of the ATAD3A hexamer due to the longer side chain. Arg327 is located next to the ATP binding pocket. The side chain of Arg327 forms a salt bridge with the carboxylate of the Glu469 side chain, and also makes a hydrogen bond interaction with the main chain carbonyl of Met470 (Figure 1D). This interaction would be abolished by the p.(Arg327Pro) variant, resulting in structural changes that could impact ATP binding. Collectively, in silico analyses suggest that these variants could impair ATAD3A function.

### *Drosophila* studies of missense variants

To investigate the functional consequences of missense variants in *ATAD3A*, we created a new *Drosophila Atad3a* (*dAtad3a*) allele based on recently developed CRISPR/Cas-9-mediated genome editing and Drosophila genetic technologies.^20-22^ To create a null allele, we introduced an artificial exon cassette carrying *attP-SA (splicing acceptor)-T2A-Gal4-polyA-attP* into the first coding intron of *dAtad3a* (referred as *dAtad3a-T2A-Gal4*) (Figure 2A). These flies produce an N-terminal portion of the dATAD3A protein as well as the Gal4 protein whose expression is under control of endogenous cis-elements of *dAtad3a*. Gal4 is a transcriptional activator that drives expression of transgenes by binding UAS (Upstream Activating Sequence) (Figure 2A). To test expression patterns of *dAtad3a*, we generated flies having a *dAtad3a-T2A-Gal4* allele with *UAS-mCD8::GFP*. We found that dATAD3A is expressed ubiquitously during embryogenesis, and that the expression pattern includes neurons in the brain and ventral nerve cord (VNC) (Figure S1). dATAD3A remains highly expressed in brain in both larval and adult stages. Moreover, dATAD3A is expressed in the adult thorax and in peripheral neurons in adult wings (Figure 2B).

**Figure 2.**
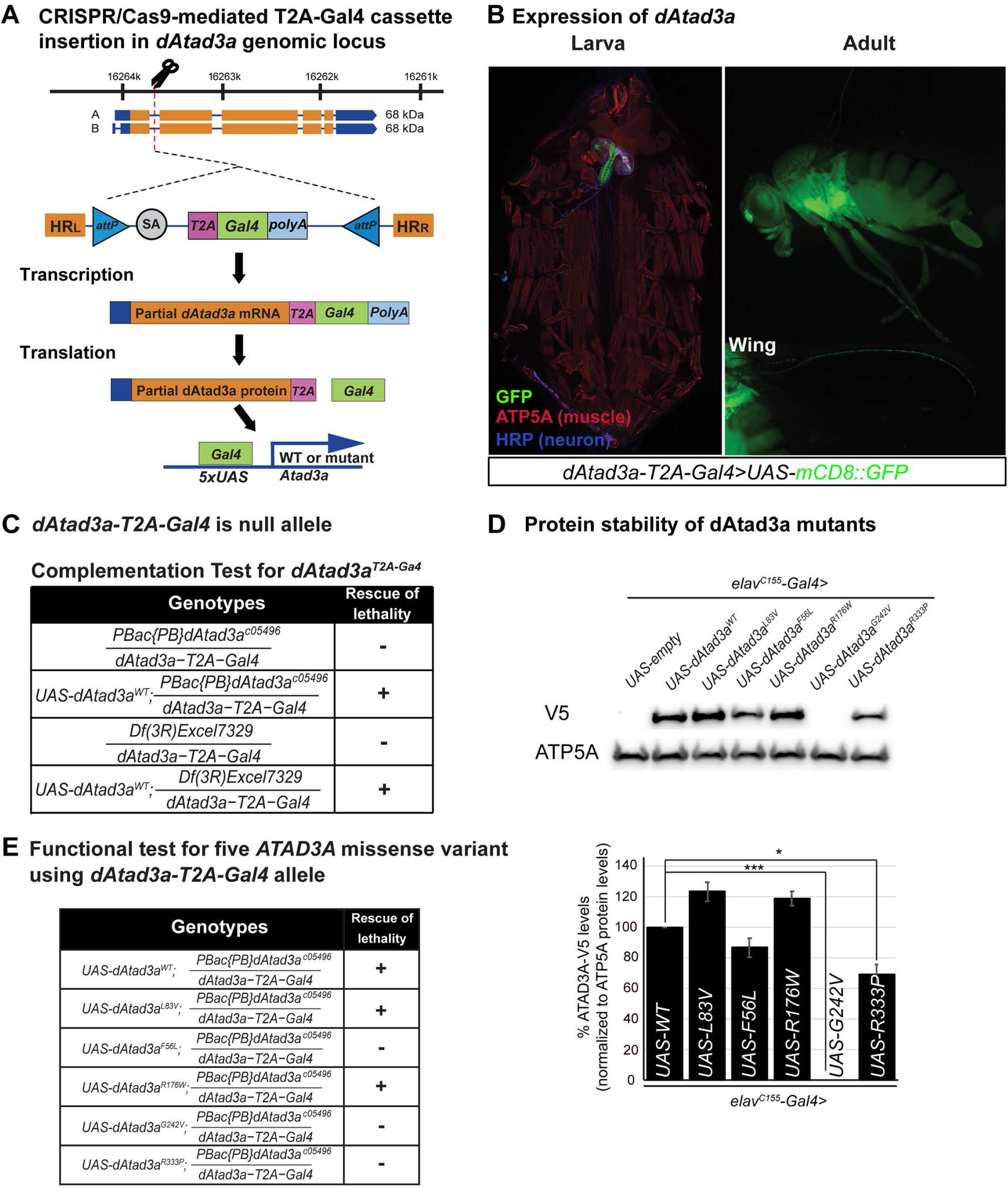
Drosophila Atad3a models shows various strength of ATAD3A missense variants. (A) A schematic of the generation of *dAtad3a-T2A-Gal4* by CRISPR-Cas9 gene editing, and the translation of a Gal4 protein by a ribosomal skipping mechanism. The location of the *attP-SA-T2A-Gal4-polyA-attP cassette* insertion into the dAtad3a genomic locus is indicated by the dotted lines. The T2A-Gal4 cassette consists of a splice acceptor (SA, light gray) followed by a ribosomal skipping T2A peptide sequence (pink), a Gal4 coding sequence (green), a polyadenylation signal (light blue). Two inverted *attB* sites (blue) are positioned at the 5’- and 3’-end of the cassette. (B) Expression of *UAS-mCD8::GFP* under the control of *dAtad3a-T2A-Gal4* is monitored in larvae and adult flies. (C) Complementation test results of *dAtad3a-T2A-Gal4* alleles. +, complement; -, failure to complement. *dAtad3a-T2A-Gal4* fails to complement a deficiency (*Df(3R)Excel7329*) that lacks the *dAtad3a* locus and *PBac{PB}dAtad3ac05496* null allele, which were rescued by expression of wildtype *dAtad3a* cDNA. These data indicate that *dAtad3a-T2A-Gal4* is a loss-of-function mutant. (D) Western blots for fly heads expressing wildtype dAtad3a-V5, or dAtad3a-V5 carrying homologous missense mutations identified from patients. Three replicates were quantified. Error bars indicate SEM. P values were calculated using Student’s t-test. *\*P* <0.05, *\*\*\*P* <0.001. (E) The lethality caused by *dAtad3a* loss was rescued by expression of wild-type *dAtad3a*, and *dAtad3a* carrying L83V, or R176W, but not by those carrying F56L, G242V, or R333P.

To test whether the *dAtad3a-T2A-Gal4* allele is a loss of function mutation, we performed complementation studies. Flies carrying a *dAtad3a-T2A-Gal4* allele and a *dAtad3a* loss-of-function allele (*PBac {PB}dAtad3a*^*c05496*^) or fly mutants lacking the entire *dAtad3a* genomic region (*Df(3R)Excel7329*) exhibited lethality in embryo stages (Figure 2C). The lethality caused by *dAtad3a* loss was fully rescued by expression of wildtype *dAtad3a* cDNA (UAS-*dAtad3a*^*WT*^) (Figure 2C). Hence, the results indicate that *dAtad3a-T2A-Gal4* is a severe loss of function allele.

To determine whether the series of missense variants in *ATAD3A* identified from affected individuals impair *in vivo* protein levels of ATAD3A, we generated transgenic flies that allow expression of *dAtad3a* cDNA carrying homologous mutations for these missense variants (*UAS-dAtad3a*^*L83V*^, *UAS-dAtad3a*^*F56L*^, *UAS-dAtad3a*^*R176W*^, *UAS-dAtad3a*^*G242V*^, *UAS-dAtad3a*^*R333P*^) under the control of UAS. We expressed each transgene together with wild-type *dAtad3a* (*UAS-dAtad3a*^*WT*^) using the pan-neuronal Gal4 driver (*elav*^*C155*^*-Gal4*). All transgenes carry C-terminal V5 tag. Western blot analysis for adult heads revealed that no protein was detected from *dAtad3a*^*G242V*^expression, and the protein levels of dAtad3a^R333P^ were lower than those in wild type control (*dAtad3a*^*WT*^) (Figure 2D). On the contrary, the protein levels from the other three transgenes (*UAS-dAtad3a*^*L83V*^, *UAS-dAtad3a*^*F56L*^, and *UAS-dAtad3a*^*R176W*^) were not significantly different than protein levels in the wild-type control (*dAtad3a*^*WT*^) (Figure 2D). Hence, these results indicate that the G242V variant is a protein null allele, and that R333P moderately affects protein levels.

To determine the effects of the series of missense variants in *ATAD3A* identified from affected individuals on *in vivo* function of ATAD3A, we tested whether expression of each missense variant rescues the developmental lethality caused by *dAtad3a* loss. We found that expression of *dAtad3a*^*F56L*^, *dAtad3a*^*G242V*^, or *dAtad3a*^*R333P*^ completely failed to rescue the lethality caused by loss of *dAtad3a*, indicating that these three variants are severe loss of function alleles (Figure 2E). Failure of lethality rescue by G242V is consistent with the Western results showing complete loss of the protein with this mutation (Figure 2D). The failure of lethality rescue by F56L and R333P could result from functional defects rather than the moderately decreased protein levels (∼20-30%) because one copy loss of *dAtad3a* (*dAtad3a*^*T2A-Gal4*^*/+* or *PBac {PB}dAtad3a*^*c05496*^*/+*) does not affect viability. On the contrary, expression of *dAtad3a*^*L83V*^ or *dAtad3a*^*R176W*^ fully rescued the developmental lethality caused by *dAtad3a* loss as we obtained adult *dAtad3a* null flies expressing *dAtad3a*^*L83V*^, or *dAtad3a*^*R176W*^ in expected Mendelian ratios (Figure 2E). In addition, *dAtad3a* mutant larvae expressing *dAtad3a*^*L83V*^, or *dAtad3a*^*R176W*^ exhibited comparable mitochondrial content to those of flies expressing *dAtad3a*^*WT*^ (Figure S2). These results indicate that the flies carrying L83V and R176W variants did not exhibit developmental defects. Hence, the results indicate that F56L, G242V, and R333P are severe loss-of-function alleles, whereas L83V and R176W only mildly affect gene function.

To investigate the phenotypic strength of F56L, G242V, and R333P, we decided to characterize phenotypes during embryogenesis from *dAtad3a* null mutants as well as each mutant because most animals expressing these variants die before the 1^st^ instar larvae stage. No reports for phenotypes caused by *dAtad3a* loss during embryogenesis have been documented so far. Biallelic deletion of *ATAD3A* and adjacent *ATAD3* paralogs in humans causes sever neuro-developmental defects including fetal congenital pontocerebellar hypoplasia and neonatal death. ^10; 12^ Thus, we sought to determine whether *dAtad3a* loss causes defects in neurodevelopment in *Drosophila* embryos using anti-Elav (a neuronal marker), and anti-HRP (a marker for neuronal membranes) antibodies. We examined stage 15 embryos in which the central nervous system (CNS) including brains and ventral nerve cord (VNC), and the peripheral neurons and their neuronal projections are well established (Figure 3A). The *dAtad3a* null mutants exhibit a wide range of neurogenesis defects. We found that loss of *dAtad3a* results in 55% of embryos with defects in CNS development and 67% with defects in PNS development (*dAtad3a* lof, Figure 3A). The CNS defects include brain mis-location, twisted and shrunken VNC, and partial absence of the VNC. The PNS phenotypes include partial absence of PNS cells, misguided PNS neural tracks, and failure of correct specification of the PNS cells. The phenotypes caused by *dAtad3a* loss were significantly rescued by expressing wildtype *dAtad3a* (*UAS-dAtad3a*^*WT*^) (29% CNS defects; 31% PNS defects), but not by expressing *dAtad3a*^*F56L*^, *dAtad3a*^*G242V*^, or *dAtad3a*^*R333P*^ (Figure 3B). Expression of F56L (48% CNS; 55% PNS), and G242V (62% CNS; 66% PNS) exhibited a phenotype strength comparable to those in null mutants, whereas R333P (60% CNS; 63% PNS) showed slightly weaker phenotypes (Figure 3B). Hence, we discovered that *dAtad3a* loss leads to severe neurodevelopmental defects in *Drosophila* embryos, and all three variants (F56L, G242V, and R333P) are severe loss of function alleles.

**Figure 3.**
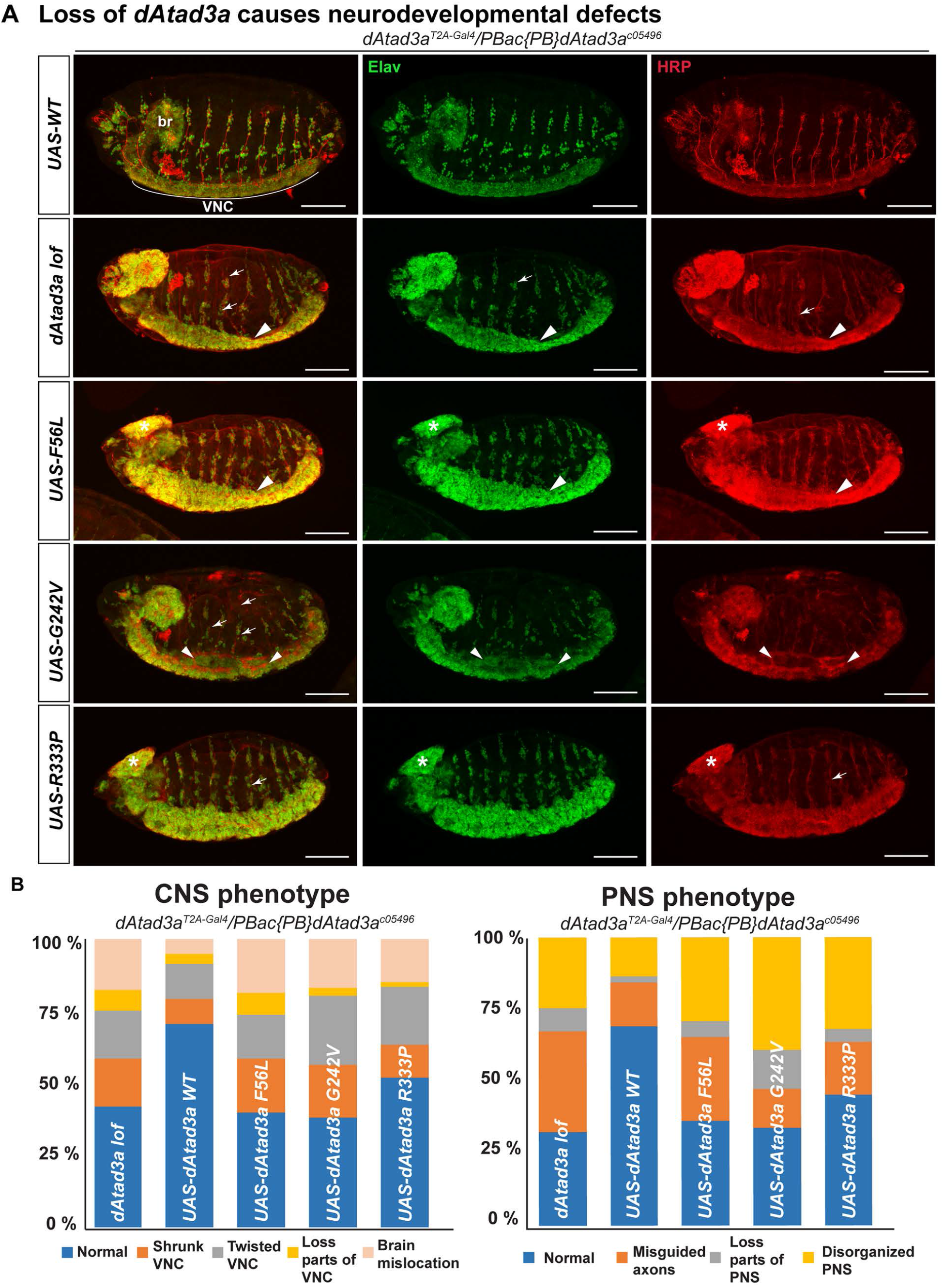
Loss of Atad3a, and F56L, G242V, and R333P variants lead to severe neurodevelopmental defects. (A) Confocal micrographs of *dAtad3a* null mutant embryos and those expressing *dAtad3a*^*WT*^, *dAtad3a*^*F56L*^, dAtad3a^G242V^, *dAtad3a*^*R333P*^. Elav (green) stained neurons, and anti-HRP (red) stained neuronal membranes. Br indicates brain, and VNC indicates ventral nerve cord. Arrowheads indicate shrunken, and twisted VNC. Arrows indicate misguided and loss of neurons in the PNS. Scale bars indicate 100 μm. (B) Quantification of CNS and PNS phenotypes shown in mutant embryos. Numbers of embryos for these analyses are as followed: CNS – wt (n=58), null (96), F56L (n=53), G242V (n=71), and R333P (n=60). PNS - wt (n=52), null (86), F56L (n=52), G242V (n=59), and R333P (n=44).

To investigate the phenotypes of L83V and R176W, we sought to characterize post-developmental phenotypes such as behavioral and age-associated phenotypes because expression of these variants did not exhibit developmental defects (Figure 2E, Figure S2). First, we performed life-span assays and found that *dAtad3a* null flies expressing L83V, or R176W exhibited shorter lifespans compared to those in flies expressing wildtype *dAtad3a* (50% survival at day 57 (R176W), day 62 (L83V), and day 64 (WT)) (Figure 4A). dATAD3A is mainly expressed in adult thorax and head (Figure 2B), thus loss of its function may affect locomotion behavior. To test this, we performed a climbing assay. We found that both variants exhibited age-dependent locomotion defects and R176W showed more severe locomotion defects compared to L83V flies (Figure 4B). We also performed a flight assay that is more sensitive than the climbing assay. Flies expressing R176W exhibited a flight defect at a young age (day 5) and failed to fly at all in old age (day35), whereas flies expressing L83V showed normal flight in young age, but mildly defective flight in old ages (Figure 4C). Collectively, these results indicate that both L83V and R176W variants are partial loss of function alleles and that R176W has more defective gene function compared to L83V.

**Figure 4.**
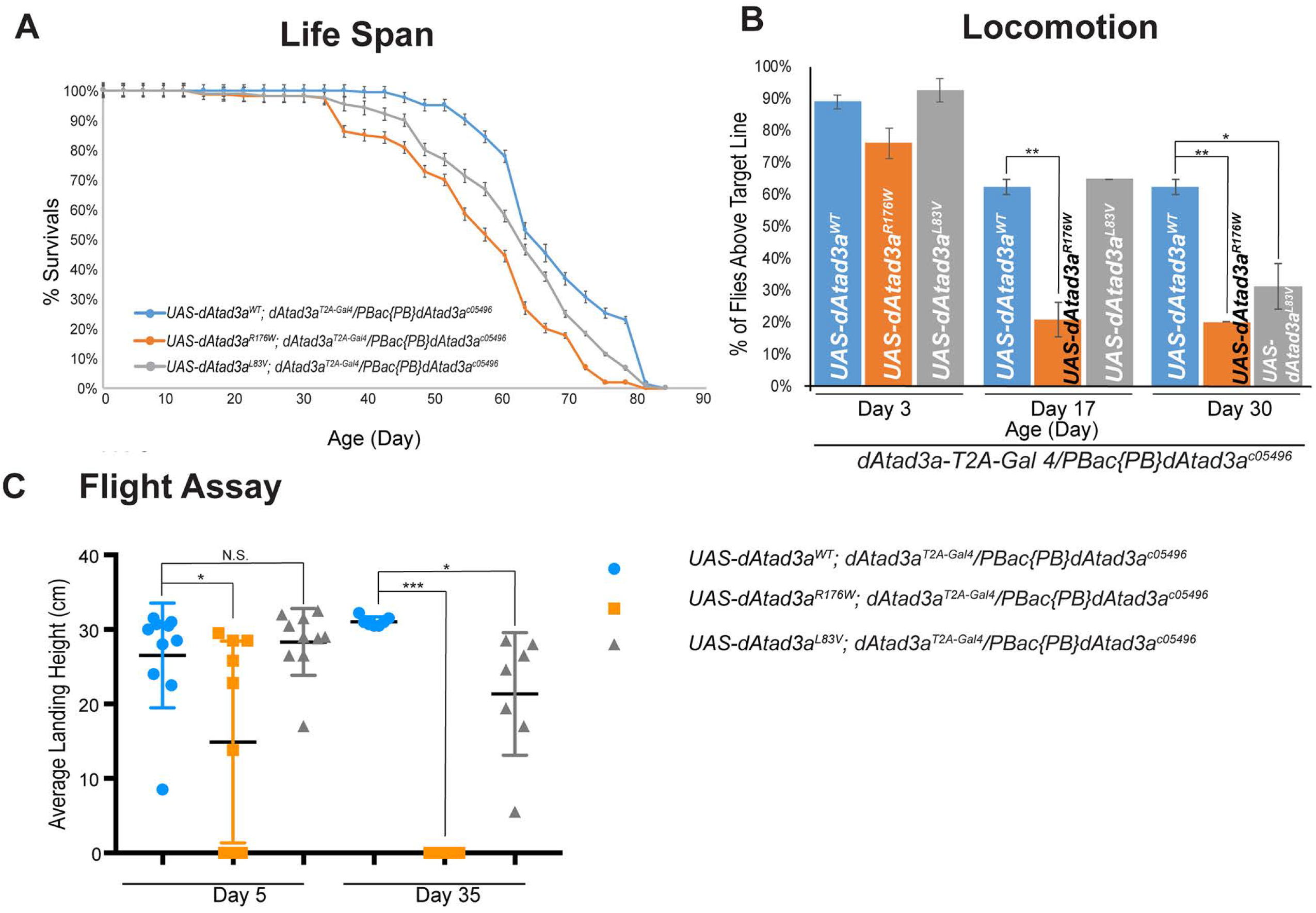
L83V and R175W variants cause behavioral defects in adult flies. (A) *dAtad3a* null mutant flies expressing L83V, and R176W were short lived compared to wildtype rescue animals. (B) *dAtad3a* null mutant flies expressing L83V, and R176W exhibited progressive climbing defects compared to wildtype rescue controls. (C) *dAtad3a* null mutant flies expressing R176W exhibited defects in flight ability in 5^th^ day of their life and complete failure of flight in 35^th^ day. *dAtad3a* mutant flies expressing L83V exhibited progressive decline of flight ability compared to rescue controls. (B, C) Three biological replicates (25 flies per group) were quantified. Error bars indicate SEM. P values were calculated using Student’s t-test. *\*P* <0.05, *\*\*\*P* <0.001.

We previously showed that human fibroblasts carrying the *de novo* variant p.Arg528Trp exhibited an increase in mitophagic vesicles and expression of *dAtad3a* carrying p.Arg534Trp, the homologous mutation of human p.Arg528Trp, leads to small mitochondria with aberrant cristae as well as an increase in autophagic vesicles in larvae muscles.^10^ These findings suggest that aberrant autophagy or mitophagy may underlie the behavioral defects in *dAtad3a* mutant flies expressing L83V or R176W (Figure 4). To test this, we sought to examine mitochondria morphology and autophagy in adult thorax muscles. First, we assessed mitochondrial morphology using an antibody for ATP5A in both young (5-day-old) and old (21-day-old) adult flies. We found that at both ages, R176W leads to smaller mitochondria with a rounded shape compared to those in wild type controls (Figure S3). On the contrary, the animals expressing L83V exhibited irregular size of mitochondria with slightly longer mitochondria on average (Figure S3). These results indicate that both R176W and L83V variants affect mitochondrial dynamics, which in turn may lead to increased autophagy or mitophagy.

To test whether *dAtad3a* carrying R176W or L83V variants cause an increase in autophagy, we measured the levels of Ref(2)P, the *Drosophila* orthologue of p62, an autophagy marker, in adult muscles. While flies expressing R176W or L83V exhibited comparable levels of Ref(2)P as compared to wild type controls as young animals (7-day-old), the older (8 week old) mutants expressing R176W or L83V, exhibited significantly higher levels of Ref(2)P than those in wild-type controls (Figure 5B). We found that in muscles expressing R176W, most mitochondria marked by ATP5A were co-localized with Ref(2)P signals. We also found large vacuole-like structures that were void of ATP5A, but Ref(2)P positive in R176W muscles (Figure 5A, arrows, dAtad3a^R176W^). In muscles expressing L83V, we found a patch of higher Ref(2)P signals that were completely void of ATP5A (Figure 5A, arrows, dAtad3a^L83V^), suggesting that mitochondria with higher Ref(2)P underwent mitophagy and were degraded through autophagosomes. To further characterize this, we performed transmission electron microscopy (TEM). TEM in 56-day-old animals revealed that R176W led to small mitochondria and increased autophagic intermediates, whereas wildtype rescue animals exhibited normal mitochondria with lower numbers of autophagic intermediates (Figure 6A and 6B). Interestingly, muscles expressing L83V showed many normal mitochondria (Figure S4) but parts of muscles were filled with autophagic intermediates (Figure 6A and 6B, Figure S4), which is consistent with the Ref(2)P results (Figure 5A). In addition, we found that R176W and L83V cause small mitochondria with bar-shape cristae (Figure 6A, 6B) as well as distinctive cristae abnormalities – cristae are loosened and torn apart (Figure S4). Collectively, the data indicate that both R176W and L83V variants lead to increased autophagy and mitochondria loss, and aberrant cristae, and that the detrimental effect of dAtad3a^R176W^ is more severe than dAtad3a^L83V^.

**Figure 5.**
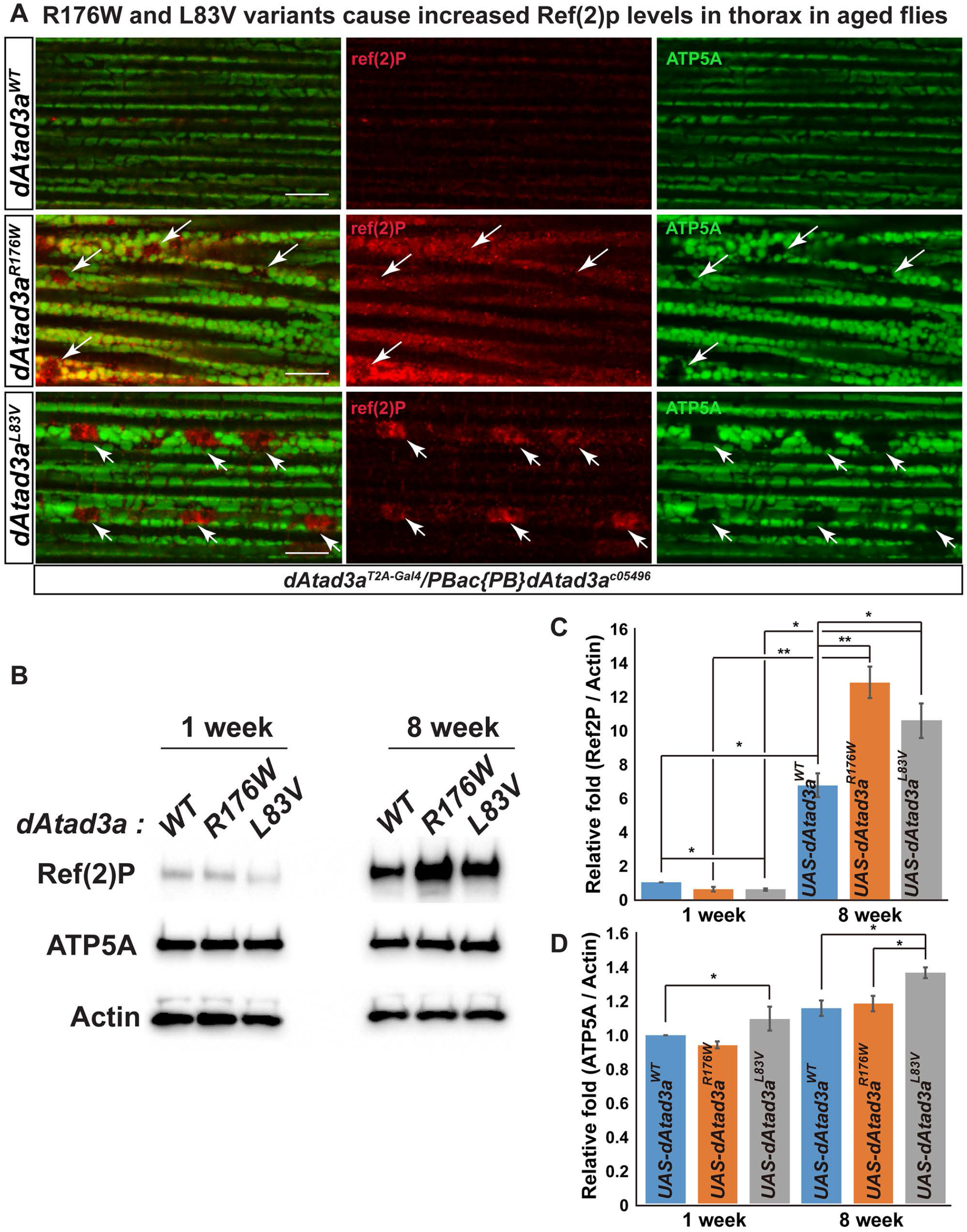
L83V and R175W variants cause increased p62 levels in thorax in aged flies. (A) Confocal micrographs of thorax muscle from 8 week-old flies - *dAtad3a* null mutants expressing *dAtad3a*^*WT*^, *dAtad3a*^*R176W*^, or *dAtad3a*^*L83V*^. ATP5A (green) labels mitochondria. Ref(2)P, is the Drosophila homolog of p62 (red). Arrows indicate Ref(2)P signals with absence of ATP5A signals. Scale bars indicate 100 μm. (B) Western blots for the protein levels of Ref(2)P, ATP5A and Actin from *dAtad3a* mutant fly thoraxes expressing *dAtad3a*^*WT*^, *dAtad3a*^*R176W*^, or *dAtad3a*^*L83V*^ (n=10 per genotype). (C) Quantification of Ref(2)P and (D) ATP5A level. Ref(2)P and ATP5A were normalized by Actin. Three biological replicates were quantified. Error bars indicate SEM. P values were calculated using Student’s t-test. *\*P* <0.05, *\*\*P* <0.01. *\*\*\*P* <0.001.

**Figure 6.**
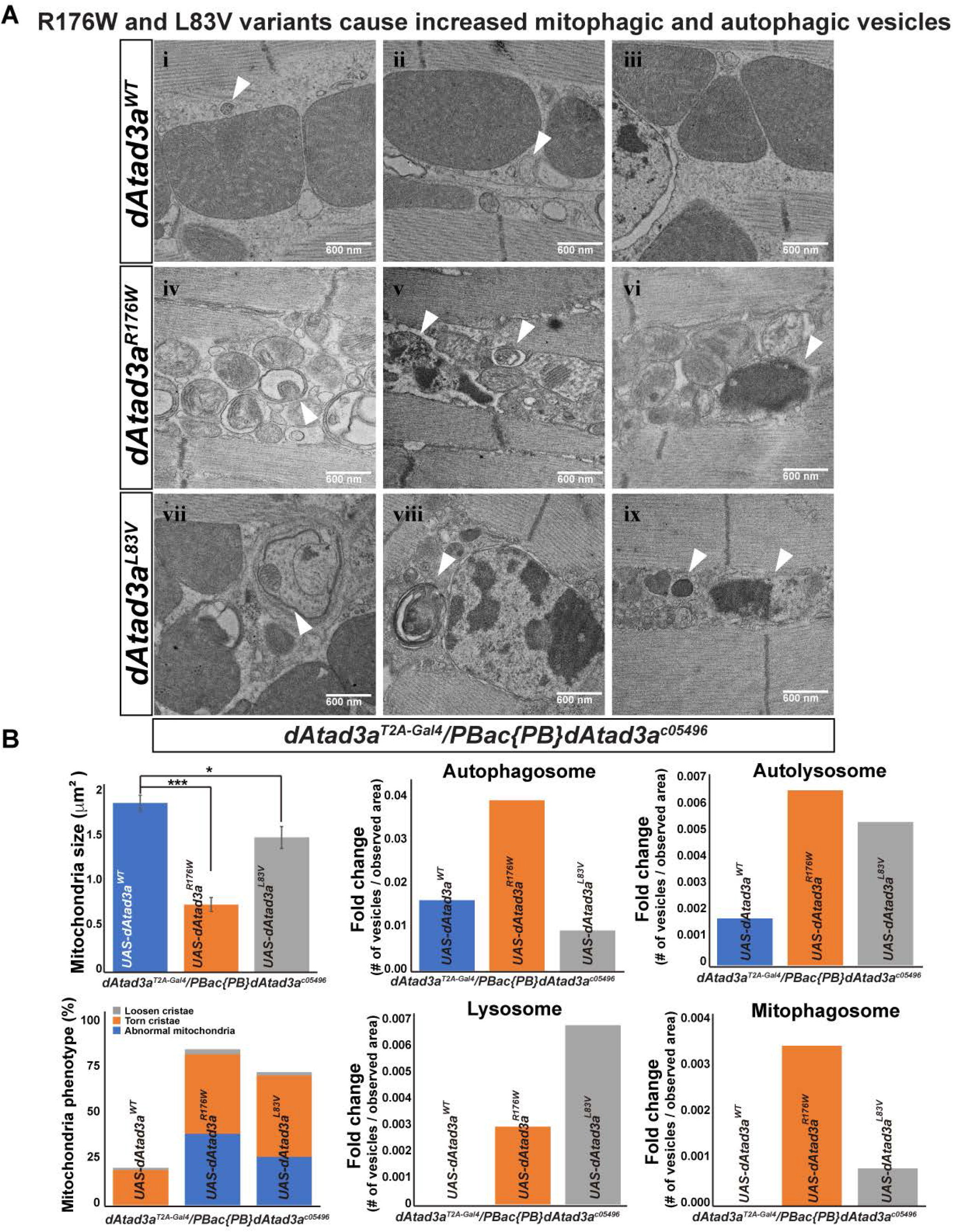
L83V and R175W variants cause aberrant mitochondrial morphology, increased autophagic and mitophagic vesicles. (A) Electron micrographs of thorax muscles from 8 week old *dAtad3a* mutant flies expressing *dAtad3a*^*WT*^, *dAtad3a*^*R176W*^, or dAtad3a^L83V^. Arrows show autophagosomes (i, ii and v), autolysosomes (v and viii), lysosomes (vi and ix), and mitophagosomes (iv and vii). Scale bars indicate 600 nm. (B) Quantification of mitochondria size, mitochondria phenotypes, numbers of autophagosome, autolysosome, lysosome and mitophagosome. Respective number of vesicles were normalized by observed area (μm^2^). Error bars indicate SEM. P values were calculated using Student’s t-test. *\*P* <0.05, *\*\*\*P* <0.001.

Functional studies for the five missense variants in *Drosophila* were consistent with bioinformatic predictions, where the human variants p.(Leu77Val) and p.(Arg170Trp) had more benign prediction scores as compared to the other missense variants (**Table 2**). However, caution must be exercised with bioinformatics prediction scores, as the p.(Phe50Leu) variant also had a relatively low conservation score (GERPrs 1.95) yet was clearly pathogenic when tested in *vivo*. Indeed, the family with the most mild phenotype, Family 5, exhibited the p.(Arg170Trp) allele in *trans* to a frameshift variant (c.1141dup; p.(Val381Glyfs*17). On the contrary, homozygosity for the p.(Arg327Pro) allele led to a severe, infantile lethal phenotype (Family 8). Genotype-phenotype correlations must take into account both alleles – the p.(Phe50Leu) allele was associated with a severe phenotype when inherited in *trans* to an intragenic deletion of two exons (Family 4); yet with a mild phenotype when observed with a nonframeshift single amino acid deletion (Family 7).

## DISCUSSION

The spectrum of *ATAD3A* variants has thus far focused on monoallelic gain-of-function variants and biallelic loss-of-function variants. We report eight families with biallelic variants, ranging from biallelic CNVs to biallelic SNVs and combinations thereof. The relatively wide phenotypic and genotypic spectrum of *ATAD3A-*associated variation calls for caution in interpretation of the clinical significance of missense variants. To address this, we systematically analyzed the functional effect of five missense variants identified in affected individuals using *Drosophila* models. We showed that F56L, G242V, and R333P are severe loss of function alleles which fail to rescue lethality of *dAtad3a* null mutants, whereas L83V and R176W are mild hypomorph variants which rescued developmental lethality, but exhibited behavioral defects in adult flies. Results from *Drosophila* studies correlated with the clinical severity of affected individuals – orthologs of the three more severe loss-of-function alleles (F56L, G242V, and R333P in *Drosophila*, or F50L, G236V, R327P in humans) led to severe phenotypes including hypotonia, global developmental delay, cataracts, cardiomyopathy and structural brain abnormalities (Families 4, 6, and 8). As noted in the results section, the F50L variant was identified in *trans* to a two exon deletion in Family 4, yielding a severe phenotype; yet was also identified in *trans* to a single amino acid deletion (nonframeshift) in Family 7, resulting in a more mild phenotype albeit with cataracts, cardiomyopathy, and cerebellar abnormalities. In contrast, one of the hypomorphic alleles (R176W in *Drosophila* or R170W in humans), when inherited in *trans* to a frameshift variant, led to a mild phenotype reminiscent of the first family reported with biallelic hypomorphic SNV.^10^ This relatively weak variant (**Table 2 - Additional file 2**) may have been overlooked or discarded in variant filtering of the exome; nonetheless, functional modeling combined with a clinical phenotype compatible with the mild range of the *ATAD3A* disease spectrum, implicates the variant as probably disease-causing.

ATAD3A belongs to the AAA+ protein family. Members of this family form oligomers and have positive cooperativity in ATP binding and hydrolysis, whereby alteration of subunits (i.e. ATPase deficient mutants) affect the function of the entire protein complex.^28^ Previous data from studies in *Drosophila* and in patient-derived fibroblasts with heterozygous variants (i.e., p.Arg528Trp; R528W and p.Gly355Asp; G355D) suggested that both alleles act in a dominant-negative fashion ^10; 14^, consistent with that the mutated residues are located in the key motifs required for the ATPase activity (Gly355, the Walker A motif^14^; R528, the Sensor 2 motif, personal communication with Sukyoung Lee). Here we show that five SNVs (missense variants) act as hypomorphic or loss-of-function alleles rather than dominant-negative. Of five variants, four variants including L77V, F50L, R170W, and G236V are located outside of the AAA+ domain (Figure 1C). Only Arg327 is located in the AAA+ domain (Figure 1D), which may impact the conformation of the AAA domain. However, human carriers for Arg327Pro (parents of family 8) are unaffected, suggesting that this allele seems not to affect the ATPase activity and does not function as dominant negative in humans. Western blot in flies showed that G236V (G242V in Drosophila) is a protein null (Figure 2D), consistent with severe loss of function of G242V *in vivo*. On the contrary, the other four alleles moderately or do not affect protein levels (Figure 2D), but rather lead to severe or partial loss of function of ATAD3A, suggesting the functional importance of the N-terminal and middle CC domains in ATAD3A. The first 50 amino acids were reported to be important to form contact sites between the mitochondria and the endoplasmic reticulum (ER) membrane^1^, implicating that F50L variant may cause defects in mitochondria-ER communication. The CC domain was shown to bind to Drp1 and oligomerization of ATAD3A leads to Drp1 to the mitochondria via the CC domain, resulting in increased fission^29^. This suggests that R170W (R176W, Drosophila), located in the CC domain, leads to small mitochondria with increased autophagosome (Figure 6B) via an increased interaction with Drp1. Further molecular studies of the pathogenic variants and identification of ATAD3A-interacting proteins will provide insight as to how genetic variants cause the etiology at the molecular and cellular levels.

Previous studies in *Drosophila* and in patient-derived fibroblasts with heterozygous variants (i.e., p.Arg528Trpand p.Gly355Asp) revealed a defect in mitochondrial dynamics, possibly triggering mitophagy and resulting in a significant reduction of mitochondria.^10^ Here we also demonstrated that aged muscles expressing R176W, and L83V variants exhibited defective mitochondrial membrane dynamics and increased mitophagic vesicles (Figure 6). Thus, these findings suggest that proper ATAD3A function is required for homeostasis of mitochondrial dynamics and mitophagy. One mechanism for mitophagy was documented in mouse hematopoietic stem cells, in which increased mitophagy in *ATAD3A-*deficient cells, has been attributed to perturbation of Pink1-mediated mitophagy.^30^ Abnormal regulation of nutrition and metabolism-sensing machineries such as mechanistic target of rapamycin (mTOR) could be implicated in the etiology caused by loss of ATAD3A as mTOR is a major regulator for autophagy and mitophagy. Indeed, Cooper et al. (2017) demonstrated upregulated basal autophagy in patient fibroblasts, associated with mTOR inactivation.^14^ In mice, *Atad3a* and mTOR have central functions in biogenesis of mitochondria during development.^2, 3, 31^ Target of rapamycin (TOR) signaling positively regulates mitochondrial activity, and the *Drosophila* paralog of *ATAD3A* (*bor*) is downregulated upon rapamycin-dependent inhibition of TOR signaling pathways.^32^ Thus, altered mTOR signaling may also contribute to the pathogenesis of *ATAD3A*-related disorders and targeting mTOR activity could be a potential therapeutic avenue for alleviating symptoms caused by *ATAD3A* mutations.

In addition to altered mitochondrial dynamics, increased mitophagy and mTOR inactivation in *ATAD3A-*deficient cells,^14, 30^ alternative pathogenetic mechanisms for *ATAD3A-* associated disorders have been proposed.^29^ These include impaired mtDNA and segregation, and aberrant cholesterol channeling and steroidogenesis.^33-35^ mtDNA co-sediments with cholesterol, and both mitochondrial integrity and cholesterol metabolism have been linked to neurodegeneration and cerebellar pathology.^12^ Fibroblasts from individuals with biallelic *ATAD3* locus deletions displayed enlarged and more numerous mitochondrial DNA (mtDNA) foci, suggesting that *ATAD3A* deficiency causes localized mtDNA aggregation or impairs its proper distribution. Moreover, fibroblasts demonstrated multiple indicators of altered cholesterol metabolism.^12^ The associated disease pathology was proposed to result either from compromised rigidity of the inner mitochondrial membrane with impaired mtDNA segregation subsequent to inadequate cholesterol metabolism, or from a shortage of cholesterol products in Purkinje cells. Affected individuals whose fibroblasts exhibited impaired cholesterol metabolism often presented with elevated urine levels of 3-methyglutaconic acid (3-MGA).^10, 12, 15^ Interestingly, *SERAC1* deficiency presents with impaired cholesterol metabolism together with elevated 3-MGA levels^37^, suggesting that defective cholesterol metabolism and mitochondrial lipid metabolism may be implicated in increased levels of 3-MGA. Whether manipulating cholesterol metabolism and mitochondrial lipid metabolism ameliorate ATAD3A pathologies remains to be investigated.

## CONCLUSIONS

*Drosophila* has been well established as a powerful genetic model organism.^36^ We utilized this model organism to assess functional impacts of various missense variants in *ATAD3A*, and showed that the allele severity in *Drosophila* correlates with the phenotypic severity in humans. We contribute to the growing disease-causing allelic spectrum at the *ATAD3A* locus, which includes biallelic NAHR-mediated deletions and a reciprocal monoalleleic duplication; monoallelic dominant-negative variants, biallelic SNVs, and now SNVs in trans to deletion alleles.

## LIST OF ABBREVIATIONS

ATAD3A: AAA-domain containing protein 3A
AAA+ ATPase: ATPases associated with diverse cellular activities
*bor*: *belphegor*
NAHR: nonallelic homologous recombination
CNV: copy number variants
SNV: single nucleotide variants
*dAtad3a*: *Drosophila Atad3a*
UAS: Upstream Activating Sequence
VNC: ventral nerve cord
CNS: central nervous system
PNS: peripheral nervous system
TEM: transmission electron microscopy

## DECLARATIONS

### Ethics approval and consent to participate

Our patients’ samples were provided for genomic analysis as an NHS diagnostic test (Family 1, 7, 8) and Technische Universietaet Muenchen (5360/12 S) (Family 2, 4, 5, and 6). The parents provided consent for genetic testing and publication of the results.

### Availability of data and materials

The datasets generated and/or analysed during the current study are available from the corresponding author on reasonable request.

## Author’s contributions

ZYY, and YP performed Drosophila experiments and provided figures. SBW, ACG, EW, KW, JAM, HL, UK, EDB, SE, and DW DSW contributed to acquisition and analysis of human genetics and clinical data. SL performed in silico structural prediction of genetic variants. LD performed EM. JRL contributed ideas for critical writing. TH and WY conceived and designed the project. TH collected and analyzed all clinical and genetic data, and wrote and revised the manuscript. WY designed and supervised the in vitro and Drosophila work, wrote and revised the manuscript. All authors edited the final manuscript.

## ACKNOWLEDGEMENTS

The authors wish to thank the families for their participation in this study. We appreciate Dr. Hugo Bellen for his generous technical support. We appreciate Dr. Scott Plafker for critical reading of this manuscript. We appreciate Dr. Sheng Zhang’s kind gift of rabbit anti-Ref2(P) antibodies.

