## Supplemental Table 1 for "Functional interpretation of *ATAD3A* variants in neuro-mitochondrial phenotypes"

**Table S1. Clinical information**

| Family number in manuscript | Family 1 | Family 2 | Family 3 | Family 4, Individual 1 | Family 4, Individual 2 | Family 5, Individual 1 | Family 5, Individual 2 |
|---|---|---|---|---|---|---|---|
| ***ATAD3A* variant 1 (NM_001170535.1)** | *ATAD3B-ATAD3A* deletion - Chr1[GRCh37):g.1425016-1462400 (maternal) | *ATAD3B* exon 4 to *ATAD3A* exon 4 Chr1:(GRCh37):g.(?_1421161)_(1453156_?)del | *ATAD3B-ATAD3A* deletion (maternal) | c.150C>G; p.(Phe50Leu) (maternal) | c.150C>G; p.(Phe50Leu) (maternal) | c.1141dup; p.(Val381Glyfs*17) (maternal) | c.1141dup; p.(Val381Glyfs*17) (maternal) |
| ***ATAD3A* variant 2 (NM_001170535.1)** | *ATAD3B-ATAD3A* deletion - Chr1[GRCh37):g.1425016-1462400 (paternal) | *ATAD3B* exon 9 *to ATAD3A* exon 9 deletion | c.229C>G; p.Leu77Val (paternal) | 3-4 exon del (c.(282+1_283-1)_(444+1_445-1))del (paternal) | 3-4 exon del (c.(282+1_283-1)_(444+1_445-1))del (paternal) | c.508C>T; p.(Arg170Trp) (not maternal; paternal DNA NA) | c.508C>T; p.(Arg170Trp) (not maternal; paternal DNA NA) |
| **Age at last exam** | 13 d (deceased due to respiratory failure) | died at 30 hours | 19 mo (died at 2yo) | died shortly after birth | died shortly after birth | 19 yr | 17 yr |
| **Gender** | male | male | female | male | male | male | female |
| **Developmental delay** | NR | NA | global DD with regression | NA | NA | mild DD; normal motor development, mod-severe learning difficulties, poor articulation | moderate DD; normal motor development, mod-severe learning difficulties |
| **Ophthalmologic involvement** | bilateral corneal dystrophy, central cataract, suspected optic atrophy | cloudy corneas | bilateral congenital cataract | bilateral cloudy corneas | NA | gaze evoked horizontal nystagmus | gaze evoked horizontal nystagmus |
| **Echocardiography** | mild LV hypertrophy (without hemodynamic relevance) | biventricular hypertrophy with poor contractility; bidirectional PDA | hypertrophic cardiomyopathy | hypertrophic cardiomyopathy | hypertrophic cardiomyopathy | global hypokinesia with LV systolic ejection fraction of 50% | thickened anterior mitral valve leaflet, no cardiomyopathy |
| **Brain MRI** | lissencephaly, hypoplastic myelin, cerebellar and midbrain hypoplasia | Post-mortem findings: microcephaly, olivopontocerebellar atrophy, bilateral ventriculomegaly, smooth parenchyma, suspected thin corpus callosum | diffuse volume loss, most severely affecting the cerebellum; apparent diffusion restriction in the hippocampal tails; normal myelination pattern | cerebellar hypoplasia, abnormal pons | cerebellar hypoplasia, abnormal pons, thin corpus callosum, ventriculomegaly, small basal ganglia with T2 hyperintensities | no cerebellar abnormality on MRI at age 20 years | mild cerebellar hemisphere volume loss, no progression on repeat MRI |
| **Seizures** | yes | yes, EEG showed burst suppression | yes, onset 20 mo | yes | NA | no | no |
| **Neurological examination** | generalized hypotonia, no respiratory drive | hypotonic from birth; no respiratory effort | hypotonia, brisk patellar DTR and positive clonus bilaterally | hypotonic from birth, no respiratory effort | hypotonic from birth, no respiratory effort | mild ataxia, impaired tandem walk, muscle wasting | wide based ataxic gait, poor tandem walk, muscle wasting |
| **Skeletal abnormalities** | multiple congenital contractures (thumbs, hips, knees, toes) | none | none | NA | NA | pectus excavatum | none |
| **Plasma lactate** | NA | elevated; CSF lactate>7.7 mmol/L | 0.6-1.9 mmol/L | NA | NA | not elevated | not elevated |
| **Metabolic studies** | tranisent hyperinsulinism | | PAA, UOA, ACP, ammonia, carnitine, VLCFA unremarkable | NA | elevataed 3-methylglutaconate in urine | NA | NA |
| **Previous genetic workup** | chromosome analysis unremarkable | CMA unremarkable | CMA unremarkable | CMA, sequencing of *EP300* and *CHD7* - unremarkable | chromosome analysis, CMA, *TMEM70* sequencing - unremarkable | CMA unremarkable | normal chromosome breakage studies, CMA revealed 1q44 deletion |
| **Other** | feeding difficulties - NG tube feeding | IUGR, undescended testes | laryngomalacia, tracheostomy dependence | cryptorchidism | NA | extreme anxiety; deafness (dx 16 years) | NA |

Abbreviations: ACP - acylcarnitine CFM - cerebral function monitor, CSF - cerebrospinal fluid, CMA - chromosomal microarray, DD - developmental delay, DTR - deep tendon reflexes, EEG - electroencephalogram, IUGR - intrauterine growth restriction, LV - left ventricle, NA - not available, NG - nasogatric, ,OAT - ornithine aminotransferase, PAA - plama amino acids, PDA - patent ductus arteriosus, PEG - percutaneous gastrostomy, UOA - urine organic acids

| Family 5, Individual 3 | Family 6 | Family 7 | Family 8, Individual 1 | Family 8, Individual 2 | Family 8, Individual 3 |
|---|---|---|---|---|---|
| c.1141dup; p.(Val381Glyfs*17) (maternal) | c.1414del; p.(His472fs) (maternal) | c.1703_1705delAGA, p.Lys568del (maternal) | c.980G>C, p.(Arg327Pro) | c.980G>C, p.(Arg327Pro) | c.980G>C, p.(Arg327Pro) |
| c.508C>T; p.(Arg170Trp) (not maternal; paternal DNA NA) | c.707G>T; p.(Gly236Val) (paternal) | c.150C>G, p.(Phe50Leu) (*de novo* ) | c.980G>C, p.(Arg327Pro) | c.980G>C, p.(Arg327Pro) | c.980G>C, p.(Arg327Pro) |
| 18 yr | 3 mo | 15 yr | 7 mo (deceased) | 6 mo (deceased) | 7 mo (deceased) |
| male | male | female | male | female | male |
| mod-severe learning difficulties | NA | mild DD, language delay, specialized school | NA | NA | NA |
| bilateral cataracts, ptosis, gyrate atrophy, nystagmus | none | bilateral congenital cataracts, intermittent strabismus divergens, mild ptosis | bilateral cataracts, vertical nystagmus | bilateral cataracts | bilateral cataracts |
| normal | hypertrophic cardiomyopathy | hypertrophic cardiomyopathy | hypertrophic cardiomyopathy, pericardial effusion | hypertrophic cardiomyopathy | hypertrophic cardiomyopathy |
| progressive cerebellar atrophy | normal | MRI (age 4 years): bilateral cerebellar lesions affecting myelination | NA | NA | NA |
| no | jitteriness and jerking, progressing to encephalopathy; EEG showed focal seizures | suspected, but EEG unremarkable | NA | NA | NA |
| unable to walk; progressive ataxia; spasticity, severe muscle wasting | central hypotonia with increased tone peripherally; severe dystonia | mild, generalized hypotonia; mild non-progressive ataxia | NA | NA | NA |
| scoliosis, Marfanoid habitus, flexion deformities at knees, metatarsus adductus | NA | scoliosis, pes valgus | NA | NA | NA |
| NA | not elevated | not elevated | NA | 3.8 mmol/L (RR 0.5-2.2) | NA |
| OAT deficiency | ammonia, UOA, ACP, PAA, VLCFA, biotindase, CSF gycine, urine AASA - unremarkable | PAA, ACP unremarkable | muscle biopsy: reduced coenzyme Q10, reduced activity complex V (30% activity) | elevated urinary methylmalonic and fumaric acid | NA |
| NA | NA | *AGK* sequencing - unremarkable | NA | NA | NA |
| deafness, feeding difficulties (PEG inserted at 14 years) | passed newborn hearing screen; right inguinal hernia and umbilical hernia | growth hormone deficiency, feeding difficulties | NA | NA | NA |
