## Supplementary figures and images for "Functional interpretation of *ATAD3A* variants in neuro-mitochondrial phenotypes"

### Supplemental Figures

Expression of dAtad3a in Drosophila embryo

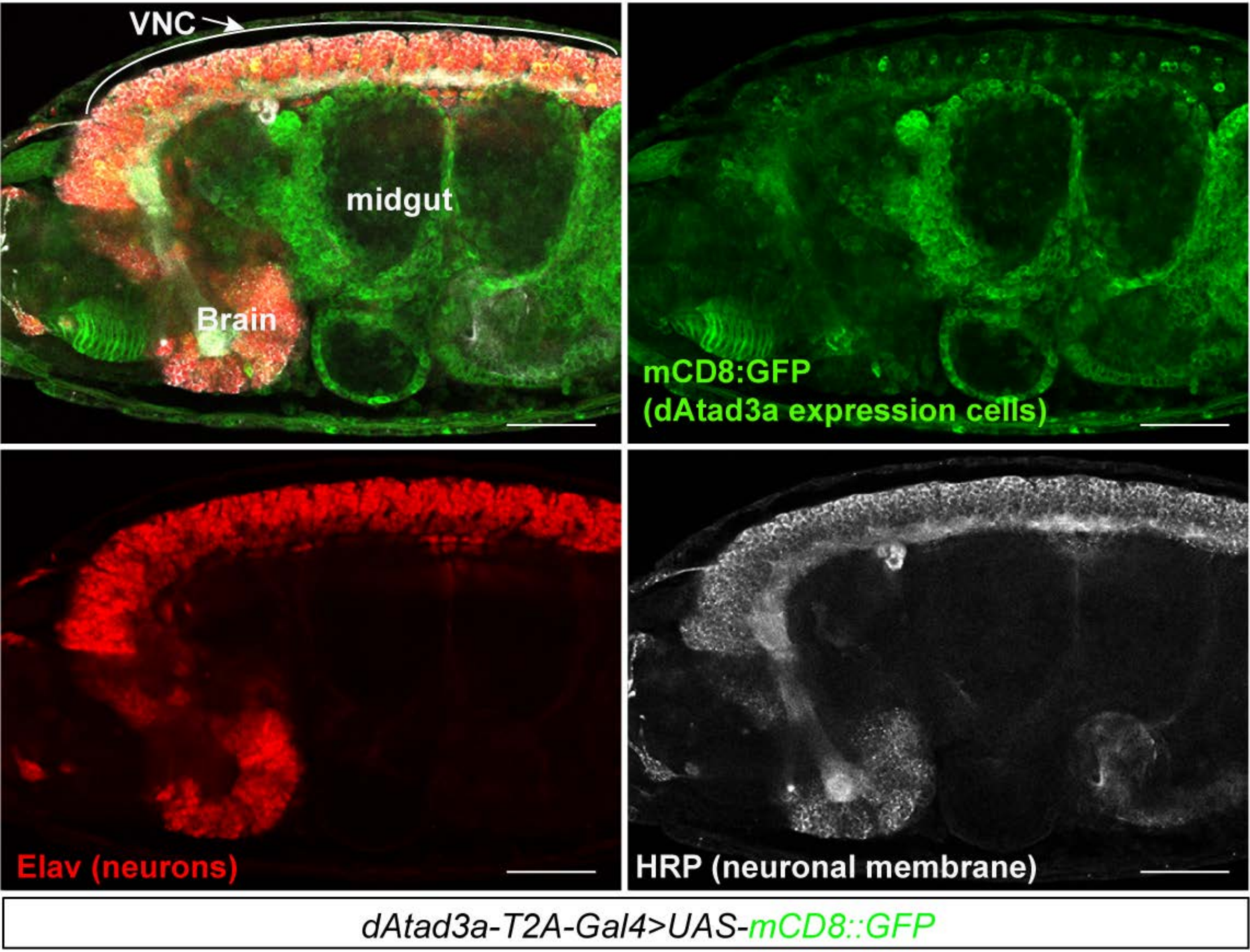


**Figure S1**

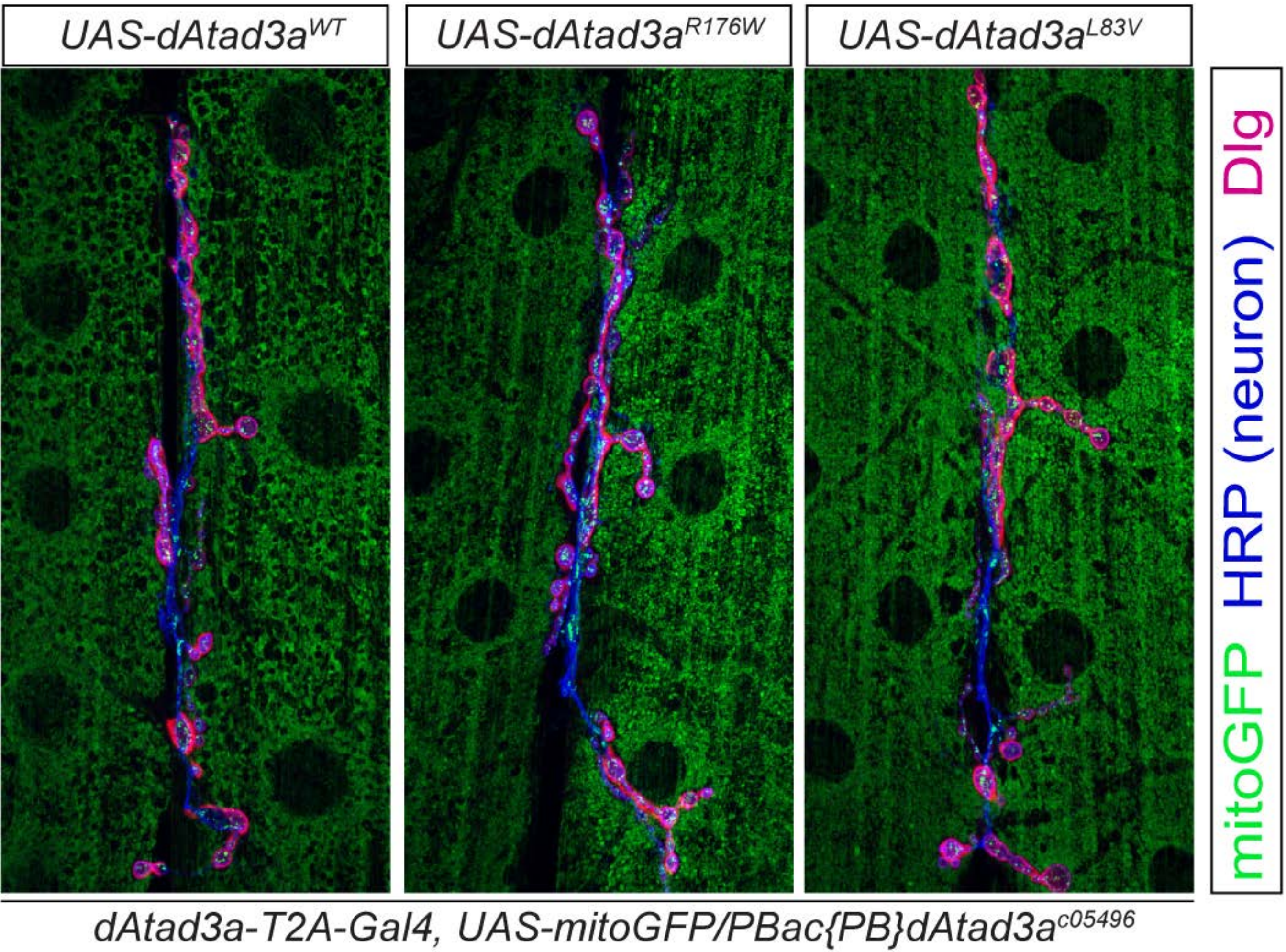

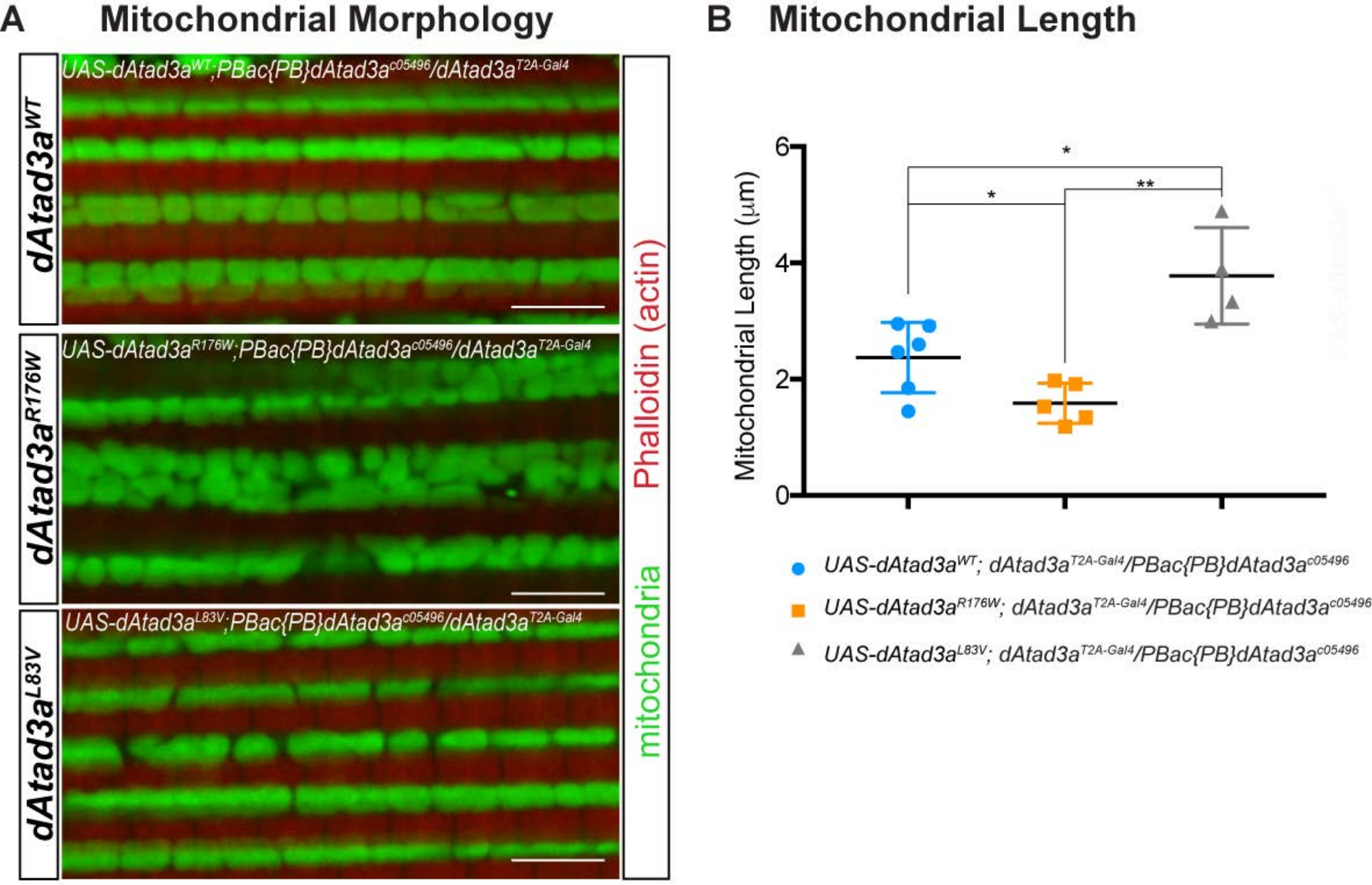

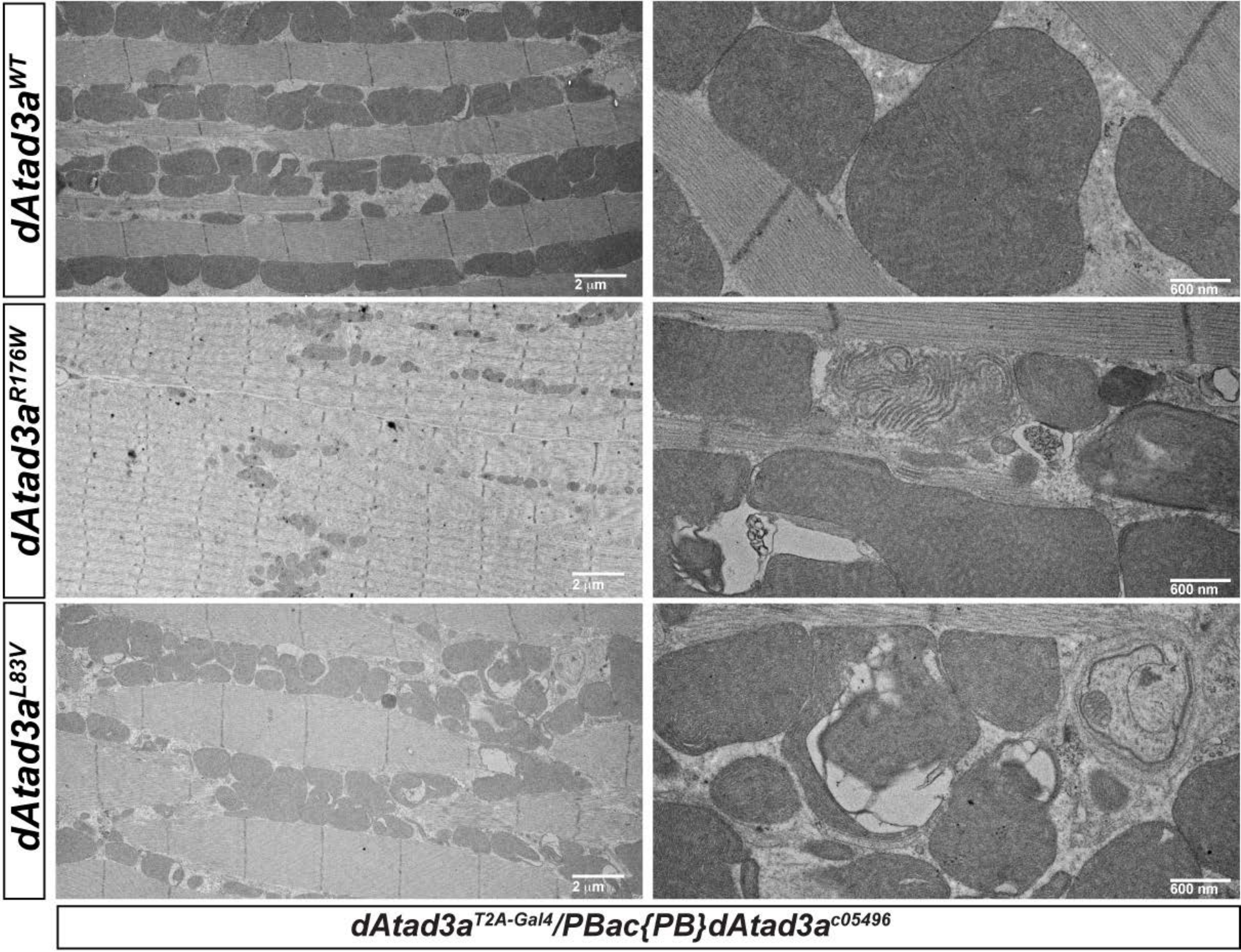
