## Supplemental Figure Legends for "Functional interpretation of *ATAD3A* variants in neuro-mitochondrial phenotypes"

**Figure S1. dAtad3a is expressed ubiquitously in embryos**

Confocal micrographs of an embryo expressing GFP protein (green) under the control of *dAtad3a-T2A-Gal4 (dAtad3a-T2A-Gal4/UAS-mCD8::GFP*). Elav (red) stained neurons. HRP stained neuronal membranes. VNC indicates ventral nerve cord. Scale bars indicate 50 μm.

**Figure S2. R176W and L83V did not affect mitochondria content and morphology in larvae muscles**

Confocal micrographs of *dAtad3a* mutant larvae muscles expressing *dAtad3a*$^{WT}$, *dAtad3a*$^{R176W}$, or *dAtad3a*$^{L83V}$. mitoGFP (green) labels mitochondria. Dlg (red) labels boutons. HRP (blue) labels neurons.

**Figure S3. R176W causes small morphology in adult muscles**

(A) Confocal micrographs of *dAtad3a* mutant adult muscles expressing *dAtad3a*$^{WT}$, *dAtad3a*$^{R176W}$, or *dAtad3a*$^{L83V}$. mitoGFP (green) labels mitochondria. Phalloidin (red) labels actin. Scale bar, 10 μm. (B) Quantification of mitochondrial length for *dAtad3a* mutant adult expressing *dAtad3a*$^{WT}$, *dAtad3a*$^{R176W}$, or *dAtad3a*$^{L83V}$. Error bars indicate SEM. P values were calculated using Student's t-test. **P* <0.05, ***P* <0.01, ****P* <0.001.

**Figure S4. R176W and L83V cause various defects in mitochondria in adult muscles**

Electron micrographs of thorax muscles from 8 week old *dAtad3a* mutant flies expressing *dAtad3a*$^{WT}$, *dAtad3a*$^{R176W}$, or dAtad3a$^{L83V}$. Scale bar, 2μm (left), and 600 nm (right).
